# Accumbens Nitrergic Interneurons Drive the Cell Type-Specific Synaptic Plasticity Required for Cue-Induced Cocaine Seeking

**DOI:** 10.1101/2024.04.05.587610

**Authors:** Adam R. Denton, Rachel E. Clarke, Lisa M. Green, Jayda Carroll-Deaton, Annaka M. Westphal, Benjamin M. Siemsen, Ashley G. Brock, Elizabeth A. Hochberg, Jacob Lavine, Makoto Taniguchi, Jacqueline F. McGinty, Alex C.W. Smith, Constanza Garcia-Keller, James M. Otis, Michael D. Scofield

## Abstract

Cocaine use disorder (CUD) remains a serious public health crisis, with relapse vulnerability continuing to pose the largest impediment to effective clinical treatment. Relapse to cocaine seeking is often triggered by drug craving evoked by exposure to drug-associated environmental cues. Data from preclinical models of rodent self-administration (SA) and cue-induced reinstatement demonstrate that exposure to drug predictive cues following a period of withdrawal engages a large induction of glutamate release in the nucleus accumbens core (NAc), not observed during cued sucrose seeking. This profound glutamate release engages neuronal nitric oxide synthase (nNOS) expressing interneurons likely through activation of metabotropic glutamate receptor 5 (mGluR5), leading to increased production of nitric oxide (NO). Importantly, cue-induced glutamate and NO production have been linked to activation of matrix metalloproteinases (MMPs) and induction of the transient synaptic plasticity in medium spiny neurons (MSNs) required for cued cocaine seeking. Recent evidence suggests that cue-induced structural and synaptic plasticity occurs predominantly in D1 Dopamine receptor expressing MSNs, yet despite these findings, how cue-induced glutamate release is translated into D1 MSN plasticity has yet to be elucidated. We show here that knockdown of nNOS is sufficient to block cue-induced reinstatement to cocaine and prevents cue-induced functional and structural synaptic adaptions specifically in D1 receptor containing MSNs. Next, we demonstrate that knockdown of mGluR5, selectively on nitrergic interneurons in the NAc, is sufficient to block both conditioned place preference (CPP) and cue-induced reinstatement to cocaine, mechanistically linking cue-associated glutamate release to NO signaling. Finally, we demonstrate that downstream of glutamate-mediated activation of mGluR5 on nitrergic interneurons and MMP activation, expression of β3 integrin receptors on D1 MSNs is required for cued cocaine seeking. Taken together, our data provide a mechanistic link between cocaine cue-induced glutamate release, activation of nitrergic interneurons and the D1 MSN plasticity required for cued cocaine seeking,

**Significance Statement:** Relapse vulnerability to cocaine is a persistent challenge to successful treatment of CUD. Relapse precipitated by drug-associated environment cues requires synaptic plasticity in MSNs. Here, we show that knockdown of nNOS, or mGluR5 on nitrergic interneurons, or β3 integrin receptors on D1 MSNs is sufficient to block cue-induced cocaine seeking. Taken together our data support the following cocaine seeking signaling cascade.

□ *Cocaine cues → PrL driven NAc glutamate release → mGluR5-mediated Activation of NAc nitrergic neurons → Activation of B3 integrins → D1 MSN plasticity → Reinstated Cocaine seeking*.

---

Cocaine use disorder (CUD) is a serious public health crisis impacting millions of people in the United States alone^1^. Relapse to cocaine use following prolonged withdrawal can be initiated by drug-predictive environmental cues and represents a serious impediment to successful treatment of CUD^2–4^. Rodent models of drug self-administration (SA), extinction (EXT) and cue-induced reinstatement (RST) of cocaine seeking are powerful preclinical tools for probing pathways and cell-types involved in cue-induced drug seeking^5^. The nucleus accumbens (NA) is a ventral striatal region critical to the integration of cortical and limbic inputs that guide motivated behavior, with the core sub-compartment (NAc) having been heavily implicated in the pathophysiology of addiction and drug seeking behaviors ^6–13^. Importantly, cortical glutamatergic input from the prelimbic cortex (PrL) has been demonstrated to be a critical driver of cue induced relapse to drug, but not natural reward seeking. ^14–16^

A substantial body of work has been completed demonstrating that cocaine SA results in functional adaptations to glutamate homeostasis in the NAc that drive cue-induced reinstatement (RST) following prolonged drug withdrawal^17^. Cocaine SA and EXT reduces basal glutamate levels in the NAc through disruption of glial glutamate homeostasis.^10,18,19^. This loss of glutamate tone engages presynaptic potentiation due to reduced activation of mGluR2/3 autoreceptors ^9,10,17^ Upon re-exposure to drug-predictive cues, spillover of glutamate into the extracellular space drives RST behavior and produces structural and functional adaptations in NAc medium spiny neurons (MSNs) associated with cue-induced drug seeking^2,6,12,13^. As mentioned above, the prelimbic cortex (PrL) delivers most of the glutamatergic input into the NAc associated with cocaine seeking, and this PrL to NAc pathway has been heavily implicated in the pathology of addiction^15,20,21^.

The NAc is predominately comprised of dopamine receptor 1 (D1) and 2 (D2) receptor expressing MSN’s with sparce populations of interneurons consisting of parvalbumin, choline acetyltransferase and nitrergic interneurons. Despite making up less than 1% of the NAc cellular composition^22^, nitrergic interneurons have previously been implicated in RST following cocaine SA^2,11^. The primary source of nitric oxide (NO) in the NAc is calcium (Ca^2+^)-dependent neuronal nitric oxide synthase (nNOS) encoded by the *Nos1* gene, which is expressed in the subpopulation of GABAergic interneurons described above, which in the NAc co-express Somatotstatin^23,24^. Previous pharmacological studies demonstrate that elevated NAc glutamate levels induced during RST stimulate metabotropic glutamate receptors (mGluRs), in particular mGluR5, which has likewise been previously implicated in RST^25–29^. We hypothesized that cocaine cue-induced glutamate buildup during RST activates mGluR5 receptors on nitrergic interneurons evoking a Ca^2+^-dependent increase in nNOS production, which in turn drives elevations in NO production. We have shown previously that elevated NO production results in activation of matrix metalloproteinases (MMPs), which is required for the structural and functional adaptions in MSNs underlying RST^2,30^. Specifically, cocaine cues promote increases in dendritic spine head diameter on MSNs, and result in transient synaptic potentiation (t-SP) including increased AMPA/NMDA ratios, a phenomenon also potentially linked to NO via the post translational modification S-nitrosylation of stargazin, resulting in increased AMPA receptor trafficking in MSNs^2,12,31^.

Glutamate release in the NAc reaches peak levels approximately 15-30 minutes into RST testing, which parallels the time course of peak cocaine seeking behavior and the associated structural plasticity and t-SP positively correlated with responding during cue-induced cocaine seeking^11^. Interestingly, it has been shown that nitric oxide release during RST closely parallels the temporal profile of cocaine cue-induced glutamate release^11^. Collectively, previous findings indicate that nitrergic interneurons and NO signaling in the NAc are critical components of the signaling cascade required for cued RST to cocaine seeking. Despite these findings, few studies to date have sought to directly manipulate nitrergic interneurons or NO signaling in the NAc in the context of cue-induced cocaine seeking. Here, we employ several small-hairpin RNA (shRNA) constructs and transgenic animals in conjunction with behavioral testing, confocal microscopy, optogenetics and electrophysiology to elucidate how NAc nitrergic interneurons translate cocaine cue-induced NAc glutamate release into the cell type-specific MSN plasticity required for cued cocaine seeking

## Materials and Methods

### Animal Subjects

#### Rats

Male and female *D1 ^Cre+^/D2 ^Cre+^* rats were bred on a Long-Evans background as part of the National Institute on Drug Abuse Transgenic Rat Project^32^ and genotyped in-house (*N=*82). During all experiments, rats were single housed and maintained in a temperature and humidity-controlled vivarium whereby food and water were available *ad libitum*. Two days prior to the start of self-administration (SA) all rats were food restricted to 20g of rat chow per day. Rats were maintained on a 12:12 hour dark/light cycle, with all experimentation occurring during the dark phase.

#### Mice

Male and female *Nos1^tm1.1(cre/ERT2)Zjh^* mice (Nos1-CreERT2, ∼25-30g) were purchased from The Jackson Laboratory (#014541, Bar Harbor, Maine MGI: 4941223) and genotyped in-house. Mice were group housed and maintained in a temperature and humidity-controlled vivarium whereby food and water were available ad libitum on a 12:12 h dark/light cycle. All animal use protocols were approved by the Institutional Animal Care and Use Committee of the Medical University of South Carolina (protocol # 01081) and were performed according to the National Institutes of Health Guide for the Care and Use of Laboratory Animals (8^th^ ed., 2011).

### Surgical Procedures

#### Rats

For all experiments, rats were anesthetized with a 4 mg/kg, intraperitoneal (i.p.) ketamine/xylazine mixture diluted in sterile saline. At the time of surgery, all rats were ∼300-400g. Rats received a chronic i.v. indwelling silastic catheter implant followed by stereotaxic injection. **In Experiment 1**, rats received a bilateral prelimbic (PrL; AP: +2.8, ML: +/-0.6, DV: -3.8) infusion of AAV2.*CamKII*::hChR2(H134R)-EYFP (Addgene, titer: ∼1×10^13^) and a bilateral infusion of either AAV2.*CMV::*nNOS-shRNA-EGFP or AAV2.*CMV::*luciferase-shRNA-EGFP combined with an AAV2.*hsyn*::DIO-mCherry (Addgene, titer: ∼7×10^12^) into the NAcore. In **Experiment 2**, All rats received bilateral microinjections of AAV2.*CMV::*nNOS-shRNA-EGFP or AAV2.*CMV::*luciferase-shRNA-EGFP (USC viral vector core: titer∼∼7×10^13^ vg/mL) combined with an AAV2.*CAG*::Flex-Ruby2sm-Flag.WPRE.SV40 (Addgene: titer: ∼1×10^12^ vg/ml) into the NAc. In **Experiment 5,** rats received a bilateral infusion of a Cre-dependent control or β3 shRNA into the NAc generated using the pSICO backbone^33^.

#### Mice

In **Experiment 3,** mice then received intra-NAcore (AP: +1.3, ML: +/- 1.3, DV: -4.4 mm relative to bregma) microinjection of either AAV5.*hSyn*::DIO-mCherry (Addgene, titer: ∼7×10^12^ viral genomes per mL (vg/mL)), AAV2.*CMV*::mCherry-U6-lox-eGFP-lox-Luciferase-shRNA, or AAV2.*CMV*::mCherry-U6-lox-eGFP-lox-Grm5-shRNA (0.4 µl/hemisphere, titers: ∼7×10^13^ vg/mL) For **Experiment 4**, mice received a chronic i.v. indwelling silastic catheter implant followed by bilateral implants of either AAV2.*CMV*::mCherry-U6-lox-eGFP-lox-Luciferase-shRNA or AAV2.*CMV*::mCherry-U6-lox-eGFP-lox-Grm5-shRNA (0.4 µl/hemisphere, titers ∼7×10^13^ vg/mL) and were allowed to recover for two weeks before experiments began. Mice in all experiments were allowed 2 full weeks of recovery prior to the start of behavioral testing.

#### Drugs

Cocaine hydrochloride was obtained from NIDA (Research Triangle Park, NC, USA) and was dissolved in sterile saline. 4-hydroxy-tamoxfen (4-OHT) was purchased from Sigma (H6728) and was dissolved in 100% DMSO and stored in the -20 until use. Immediately prior to administration, frozen aliquots were diluted in sterile saline containing 2% Tween-80 for a final concentration of 2 mg/ml 4-OHT in 5% DMSO. 4-OHT was administered to mice at 20 mg/kg intraperitoneally. Animals were maintained in the colony for 4 weeks post-4-OHT administration until experiments began.

#### Rat Cocaine SA (Experiments 1,2 and 5)

Following recovery, rats were food restricted to 20 g of chow per day for the duration off all experimentation. Rats received non-contingent saline infusions (Yoked Saline; YS) or were trained to self-administer cocaine on an FR 1 schedule of reinforcement during 2 h sessions for 10-12 days in operant boxes contained inside sound-attenuating ventilated chambers (Med Associates, Fairfax, VT, USA). Active lever presses (ALP) elicited a light plus tone conditioned cue followed by a single infusion of cocaine hydrochloride (NIDA, Research Triangle Park, NC, USA; 200 μg/50 μl bolus) followed by a 20s time out period whereby ALP elicited no consequence to prevent rodent overdose. Inactive lever presses (ILP) elicited no cues or reward delivery. Following acquisition of SA, rats underwent extinction training (EXT) for a minimum of 5 days until extinction criteria was met (::25 ALP over last 2 days of extinction). Animals were returned to the vivarium overnight before undergoing 15 minute cued-reinstatement testing (RST) the following morning. 15 minute RST testing was performed as peak glutamate and NO concentrations during cue-induced drug seeking occur approximately 15 m into the seeking session^11^. During RST, ALP resulted in the presentation of both light and tone cues, but no cocaine delivery, whereas ILP had no consequence. All behavior data were extracted in 5 m bins through MedPC software.

#### Mouse Conditioned Place Preference (Experiment 3)

Conditioned Place Preference to cocaine (7.5 mg/kg) was conducted as previously described ^34^. During the pre-test, mice were allowed to freely explore both chambers of a 2-chamber conditioned place preference box (Med Associates) for 20 minutes prior to conditioning days. Animals were counterbalanced based on pre-test preference to assign cocaine-paired chamber. Animals received 4 pairing sessions (2 saline and 2 cocaine (7.5 mg/kg), where animals received saline (morning) or cocaine (afternoon) i.p injections and were isolated to pairing chamber for 30 minutes. On the test day, animals were allowed to freely explore both chambers for 20 minutes and the time spent in each chamber was recorded. Data is expressed as time spent in the cocaine paired side – time spent in saline paired side during the test.

#### Mouse Cocaine Self-Administration (SA, Experiment 4)

Mice were trained to self-administer cocaine on a fixed ratio (FR1) schedule of reinforcement in standard mouse MedPC operant chambers^2,16^. I**nactive lever pressing (ILP)** had no consequence while **active lever pressing (ALP)** resulted in the presentation of a light/tone cue plus i.v. cocaine infusion (∼1mg/kg/infusion). Mice were trained 5 days/week across 10-15 sessions until meeting criteria (at least 3 days of 15 cocaine infusions). Following SA, mice received at least 6 days of EXT training, where levers presses have no consequence. Criteria for extinction was set at <25 ALP across two EXT sessions. Once extinguished, mice underwent cued reinstatement testing, where ALP resulted in both the light + tone cue without the delivery of cocaine or sucrose.

#### Transcardial Perfusions and Immunohistochemistry (Experiments 1, 4 and 5)

Immediately following RST testing, mice and rats were heavily anesthetized with ∼1.0-2.0 ml of urethane (Fisher Scientific, CA#AC325540500, 30% w/v, i.p.) and transcardially perfused with 120 ml of 1% PB followed by 180 ml of 4% granular PFA in PB. Brains were extracted, post-fixed for 24 h and 80μm coronal sections containing the NAcore were collected using a vibrating microtome (Leica Biosystems Inc, Deer Park, IL, USA). For structural plasticity analyses (**Experiment 2**), free-floating tissue sections were then blocked in 0.1M PBS with 2% Triton X-100 (PBST) and 2% goat serum (NGS, Jackson Immuno Research, Westgrove, PA, USA) for 2 hours with agitation. Sections were then incubated overnight with agitation at 4°C with mouse anti-Flag (Sigma-Aldrich Cat# F1404, RRID: AB_262044) diluted in 2% PBST and 2% NGS, before being washed 3 times for 5 minutes in PBST. Tissue sections were then incubated in the appropriate secondary antibody conjugated to Alexa Fluor 594 diluted in 2% PBST and 2% NGS at 2-4 hours room temperature, covered, with agitation, before being washed 3 times for 5 minutes in PBST. Tissue sections were then mounted with ProLong Gold antifade (ThermoFisher Scientific, Waltham, MA, USA) and coverslipped. Slides were stored at 4°C protected from light until imaging. To verify the extent of nNOS knockdown, sections from a subset of animals (n=16) were processed identically as described above with rabbit anti-nNOS (Millipore Cat# AB5380, RRID AB_91824) and the appropriate secondary antibody conjugated to Alexa Fluor 647. For all experiments, all primary and secondary antibodies were used at a concentration of 1:1000. All YS receiving animals were sacrificed 24 h following the final EXT session.

#### Patch-Clamp Electrophysiology (Experiment 1)

Rats were anesthetized with isoflurane and rapidly decapitated for brain extraction either 24 h following the final extinction session (YS) or immediately following RST (cocaine SA). Brains were rapidly removed and bathed in ice-cold, oxygenated (95% O2, 5% CO2; Airgas) sucrose-based cutting solution containing (in mM): 225 sucrose, 119 NaCl, 1.0 NaH2PO4, 4.9 MgCl2, 0.1 CaCl2, 26.2 NaHCO3, 1.25 glucose (305-310 mOsm). Coronal sections 300 μm thick were taken using a vibratome (Leica VT1200) and immediately transferred to oxygenated, warm (32◦C) aCSF containing the following (in mM): 119 NaCl, 2.5 KCl, 1.0 NaH2PO4, 1.3 MgCl, 2.5 CaCl2, 26.2 NaHCO3, 15 glucose (305-310 mOsm).

Following at least 30 min of recovery, slices were constantly perfused with oxygenated, room-temperature aCSF containing picrotoxin (100 uM; 1 mL/min). NAcore cells were visualized using differential interference contrast (DIC) through a 40x liquid-immersion objective mounted on an upright microscope (Olympus BX51), and a microscope-mounted camera (Scientifica, SciCam Pro, Oculus). NAcore MSNs expressing mCherry were visualized using an integrated green LED (545 nm; <1 mW) and a GFP epifluorescence filter set. Whole-cell recordings were obtained using borosilicate pipettes (3-7 Mohm) back-filled with a potassium gluconate-based internal solution composed of the following (in mM): 135 K-Gluconate, 10 HEPES, 4 KCl, 4 MgATP, 0.3 NaGTP (pH 7.35, 280 mOsm) to characterize intrinsic excitability and spontaneous synaptic currents. Alternatively, recordings were obtained using a cesium methanesulfonate-based internal solution composed of the following (in mM): 117 Cs-Methanesulfonate, 20 HEPES, 0.4 EGTA, 2.8 NaCl, 5 TEA, 4.92 MgATP, 0.47 NaGTP (pH 7.35, 280 mOsm) to measure the amplitude of synaptic currents (mediated by AMPA and NMDA receptors). Electrophysiological data acquisition occurred at a 10-kHz sampling rate through a MultiClamp 700B amplifier connected to a Digidata 1550B digitizer (Molecular Devices) and were analyzed using Clampfit 11.2 (Molecular Devices).

Current-clamp recordings were obtained from mCherry+ and neighboring mCherry-NAcore MSNs to characterize intrinsic excitability. During recordings, resting membrane potential (Vm) was recorded and cells were polarized to -70 mV to adjust for resting membrane potential differences between cells for the remainder of the recordings. Rheobase data were collected by applying a series of 50 ms sweeps at +10 pA steps starting at 0 pA until the neuron fired a single action potential. Action potential firing was characterized by applying 500 ms depolarization sweeps at +50 pA steps from -100 to 400 pA. Spontaneous excitatory postsynaptic currents (sEPSCs) were recorded in voltage-clamp mode over a 2 min period.

Voltage-clamp recordings were obtained from eGFP+ (shRNA construct) and mCherry+ and neighboring mCherry-NAcore MSNs to characterize inputs from PrL. During recordings, NAcore MSNs were held at -70 mV and PrL axons containing AAV2-EF1a-DIO-hChR2-EYFP were activated through a 10 ms pulse of a blue LED (470 nm; 1 mW) applied every 5 s, resulting in optically evoked excitatory postsynaptic currents (oeEPSCs). AMPAr/NMDAr ratios were collected by optogenetically evoked EPSCs at holding potentials of -80mV and +50mV to isolate fast AMPAr- and slow NMDAr-mediated oeEPSCs, respectively. AMPAr/NMDAr ratios were calculated by dividing the amplitude of the AMPAr current (peak current, at -80 mV) by the amplitude of the NMDAr current (current at 50 ms following start of pulse, at +50mV). While AMPAr oeEPSCs and AMPAr/NMDAr ratios were generally recorded using Cs-methanesulfonate, in a subset of cells we recorded AMPAr oeEPSCs using potassium gluconate internal solution.

### RNAScope™ (Experiment 3)

Mice were rapidly decapitated and fresh frozen brains were sectioned on a cryostat (Leica instruments) at 14 µm and adhered to slides and stored at -80°C for ≤3 months before performing RNAScope™. The RNAScope™ Multiplex Fluorescent v2 Kit (ACD bio, Newark, CA) was used according to the manufacturer’s instructions with minor adjustments. Following fixation and dehydration, endogenous GFP and mCherry proteins were removed using the protease IV reagent for 10 minutes. The following probes for mRNA detection were used: Mm-Grm5 (C1, #423631), Mm-Nos1 (C2, #437651), and EGFP (C3, #400281). mRNA probes were detected using TSA®Plus detection kit (Akoya Biosciences, Marlborough, MA); eGFP was detected with Fluoroscein (1:2000), Nos1 with Cy3 (1:2000), and Grm5 with Cy5 (1:1500). Slides were stored at 4°C protected from light until imaging.

#### Mapping of virus expression

All imaging was performed by an investigator blind to experimental conditions. To determine accuracy of microinjections into the NAc, a Leica TCS SP8 (Leica Microsystems, Deer Park, IL, USA) was used to generate low-magnification tile scans of coronal tissue sections. Expression profiles were used to generate heatmaps of approximate viral spread and placement accuracy.

#### Confocal microscopy and analyses: Experiment 2

Coronal sections containing the NAc were processed for Flag (neuronal label), EGFP (shRNA expression) and nNOS (shRNA verification) and imaged. NAcore neurons were imaged with an OPSL 552 nm laser line while nNOS was imaged with a Diode 638 laser line. Dendrites were selected for imaging based on the following criteria: (1) relative isolation from interfering dendrites; (2) location past the second branch point from the soma; (3) traceability back to the soma of origin and (4) location within the field of nNOS shRNA expression indicated by unamplified EGFP expression. Images were acquired using a 63X oil-immersion objective (1.4 N.A.) with a frame size of 1024 X 512 pixels, a step size of 0.1 μm, 4.1X digital zoom and either a 1.2 AU pinhole (Flag) or a 1.0 AU pinhole (nNOS). Laser power and gain were first optimized and held relatively constant with minimal adjustments used only to maintain voxel saturation consistency between images. Images were deconvolved using Huygens software (Scientific Volume Imaging, Hillversum, Netherlands) prior to analysis^20^. Deconvolved Z-stacks were imported to Bitplane Imaris (v9.0, Zurich, Switzerland). The filament module was used to semi-manually trace identified dendritic spines^16,20,35^. Dendritic spine head diameter (dH) was calculated using an automatic threshold set by Imaris, while dendritic spine density was calculated by normalizing the number of spines to the length of the dendrite (in μm). All analyses were carried out by an investigator blind to treatment conditions. For quantification of the extent of shRNA induced knockdown of nNOS expression, images were acquired with a 10X objective with a frame size of 1024 X 1024 pixels, 2.41 μm step size, 1.0 AU pinhole (EGFP) or a 1.5 AU pinhole (nNOS). Non-deconvolved Z-stacks were imported into Imaris, and the spots module was used to semi-manually label nNOS+ cells in the region of unamplified EGFP (shRNA) expression. **Experiment 3:** For validation of knockdown efficacy of mGluR5 shRNA, all imaging parameters for each experiment were empirically determined; laser power and gain were adjusted only to avoid regions of dense saturation whereas all other parameters (zoom, frame size, pinhole, step size) were held constant to ensure optical section thickness did not drift across images. RNAScope™ images (voxel size: 90x90x500 nm voxel size) were acquired with a 63X (1.4 N.A.) oil-immersion objective and were acquired such that an EGFP+ cell was in the same frame juxtaposed to a Nos1+ cell to increase the probability that each Nos+ cell analyzed was indeed virally transduced. 7-11 cells were imaged per animal from a subset of Luciferase-shRNA and Grm5-shRNA mice. Imaged z-stacks were next imported into Imaris, where Z-stacks were 3D cropped to the border of the Nos1+ cell. Next, a course surface (grain size=1 µm) was built to isolate the volume occupied by Nos1 mRNA. The Grm5 signal contained within the volume occupied was then 3D rendered as a surface. Surface thresholds were typically automatically determined, however in instances whereby the surface algorithm did not accurately label Nos1 or Grm5 mRNA the threshold was manually adjusted to best match the signal by an experimenter blind to conditions. The volume (in µm^3^) of the Grm5 signal was then normalized to the volume of the Nos1 signal and expressed as a Grm5 volume/Nos1 volume ratio.

### Statistical analyses

All statistical analyses were performed using IBM SPSS (v29 Somer, NY, USA) and GraphPad Prism (v10.0.2, La Jolla, CA, USA). Validation of novel viral vector constructs was performed using nested t-tests (control vs. shRNA). For comparisons of SA and EXT data, mixed model ANOVAs were employed to examine between subjects’ main effects (e.g. genotype, sex, viral infusion) and within-subjects’ effects of lever pressing (e.g. ILP, ALP) and all associated interaction effects. When comparing reinstatement data across all experiments, mixed model ANOVAs were employed to examine between subjects’ main effects and within-subjects’ effect of drug reinstatement (Average ALP last 2 extinction days vs. cued-reinstatement ALP) and all associated interaction effects. Ephys data were collapsed across biological sex and compared with nested ANOVAs. Spine head diameter data were compared with nested ANOVAs to examine main effects of treatment group and associated interaction effects. Frequency distributions of spine head diameter were compared with a Chi square test of independence to evaluate global distributional shifts and were followed up with a Z-test of proportions to examine within bin differences between treatment groups. Conditioned Place Preference data were compared with an independent samples t-test with a Welch’s correction for violations of homoscedasticity. All reported p-values are exact unless p<0.001. All graphs were produced using GraphPad (10.02).

## Results

### Experiment 1-2: Validation of Novel nNOS shRNA

To test the hypothesis that NAc nNOS is required for cued cocaine seeking and D1 vs. D2 MSN plasticity, we developed a novel shRNA construct driven under the expression of the CMV promoter that expresses both eGFP and a luciferase (control) or NOS1-specific shRNA. **(See Virus Schematic - Figure 1A).** We evaluated the extent of shRNA-mediated nNOS knockdown using tissue sections from a subset of animals in Experiment 1 (*N=16,* 3 sections per animal). Sections were stained with rabbit anti-nNOS and imaged. The “spots” tool in Bitplane Imaris was used to detect nNOS+ cells within the field of eGFP expression (viral vector transduction indicator). A nested t-test was used to evaluate the extent of nNOS knockdown. Compared with luciferase (control) shRNA treated animals, nNOS shRNA treated animals exhibited an approximate 80% elimination of nNOS+ cells within the field of eGFP expression [t(14)=14.21, p<0.001; **Figure 1B**; data are expressed as % of control condition]. Representative images demonstrating selective elimination of nNOS expression within the field of of eGFP expression (viral transduction) are displayed in **Figure 1C**.

**Figure 1:**
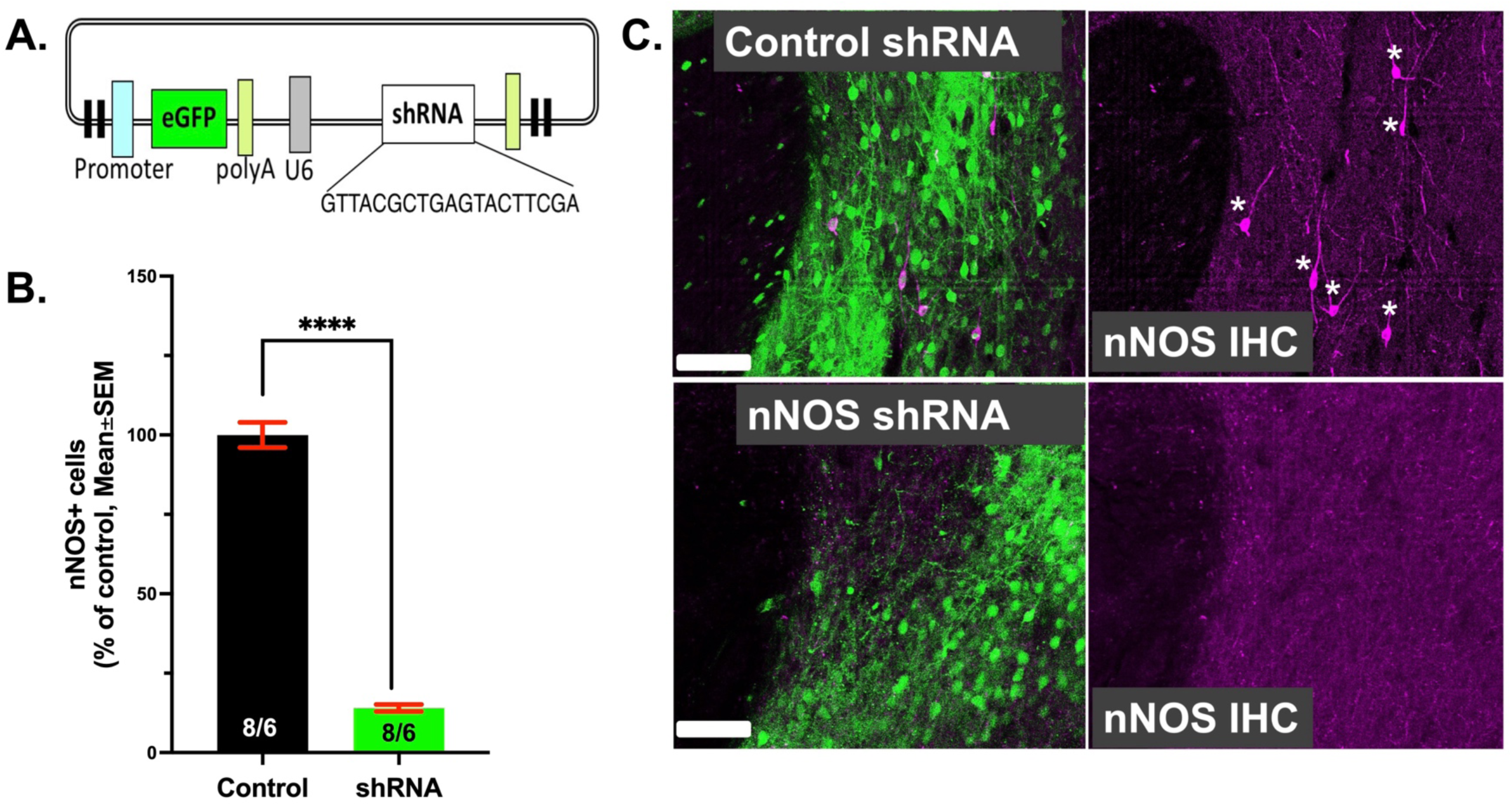
Validation of Novel nNOS shRNA: **A.)** Schematic of novel nNOS shRNA construct. **B.)** IHC validation of nNOS knockdown in the NAc. shRNA infusion resulting in an approximate 80% removal of nNOS expression in the NAc. **C.)** Representative images for control shRNA and nNOS shRNA showing field of eGFP expression and merged nNOS+ cell expression (left) in addition to isolated nNOS+ cell expression (right). *Scale bar= 50 μm; ****p<0.001*.

### Experiment 1-2: Knockdown of nNOS Blunts RST to Cocaine Seeking

To evaluate the contribution of nNOS to cocaine RST following EXT training, rats received a bilateral infusion of either control (luciferase) shRNA or nNOS shRNA into the NAc (**Figure 2A**) before undergoing cocaine SA and EXT training and RST trials. Within our study we also included yoked saline controls (YS; **Figure 2B**). No genotype, sex, or viral treatment group differences were detected during SA or EXT; data are expressed as YS verses cocaine receiving animals (**Figure 2C**). Among animals trained to self-administer cocaine, ALP was statistically significantly higher across all SA days, indicating acquisition of SA behavior [F(1,21)=61.81, p<0.001]. Upon 15 minutes RST testing, a time point shown previously to contain the largest extent of cocaine cue-induced structural plasticity^6^ animals treated with nNOS shRNA displayed significantly reduced ALP compared to control shRNA receiving animals, indicating that loss of nNOS was sufficient to blunt cue-induced reinstatement to cocaine [**Figure 2D**; F(1,22)=6.59, p=0.018]. In Figure 2D, 15 minutes of cocaine seeking data is compared to Ext responding as these animals were used in the experiments described below.

**Figure 2:**
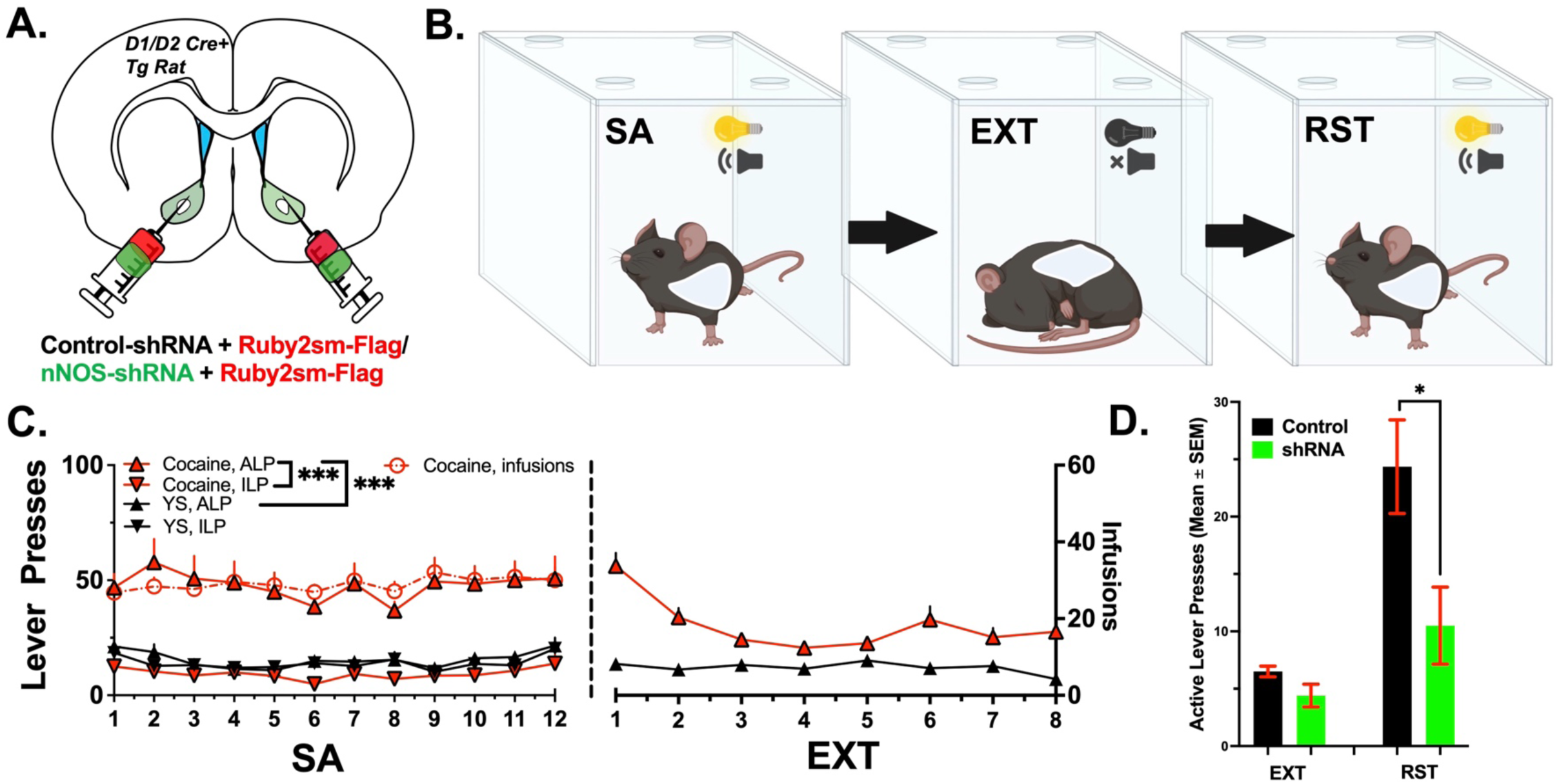
nNOS shRNA Blocks Cued Relapse to Cocaine. **A.)** Schematic indicating viral design. **B.)** Schematic indicating experimental design for behavioral experiments. **C.)** SA and EXT data for all animals. Data are collapsed across viral treatment (nNOS vs. control shRNA) as nNOS shRNA was not found to alter SA or EXT behavior. Cocaine treated animals demonstrated increased responding on the active lever, relative to the inactive lever, indicating acquisition of behavior. Cocaine treated animals demonstrated increased responding on the active lever relative to YS treated animals, indicating motivated lever pressing behavior during SA trials. **D.)** Cued reinstatement data for control and shRNA treated animals. nNOS shRNA significantly reduced active lever responding during cued reinstatement testing. No genotype or sex differences were found during cued reinstatement testing. *\*p<0.05; ***p<0.01*.

### Experiment 1: nNOS is Required for intrinsic and optically evoked D1 MSN Plasticity

In a subset of cocaine naïve animals, we provided dual injection of cre-dependent (DIO) mCherry and either luciferase (control) shRNA or nNOS shRNA, to test the hypothesis that nNOS is required for plasticity in D1 MSNs **(Figure 3A)**. All animals were yoked saline (YS) and received programmed saline infusions at intervals matched to cocaine SA trials from the behavioral experiments described above. Animals were sacrificed 24h following a final extinction session. Metrics of intrinsic excitability were recorded from eGFP+ and mCherry+ MSNs, which were either *D1+* or *D2+,* depending on rodent genotype. Metrics were likewise recorded from neighboring eGFP + and mCherry-MSNs, indicating *putative D1+* or *D2+* MSNs, allowing for simultaneous recordings of both cell types of MSNs within each animal, specifically within shRNA virally transduced neurons. Separate nested ANOVAs were used to evaluate effects of viral treatment and sex in D1 and D2 MSN populations. nNOS shRNA significantly reduced the average maximum spike number across increasing steps of injected current (**Figure 3B)** for both D1 and D2 MSNs [F(1,91)=15.15 *p<0.001*] and resulted in a reduction of average maximum evoked spikes for D1 MSNs treated with nNOS shRNA [F(1,23.31)=10.95, *p=0.003*]. Though there was a similar reduction in maximum evoked spikes for D2 MSNs, the effect was not statistically significant (**Figure 3C**). No sex differences were found for either metric. To recapitulate cocaine cue-induced PrL-derived glutamate release in the NAc, in the same animals we infused excitatory opsin in the PrL and stimulated PrL terminals in the NAc, while recording AMPAr/NMDAr ratios from virally labeled NAc MSNs (**Figure 3A).** There was a likewise reduction in AMPAr/NMDAr ratios for D1 [F(1,4.25)= 8.106, *p=0.04*), but not D2 MSNs upon optical stimulation of PrL terminals in the NAc (**Figure 3D**). Collectively, these results indicate that nNOS is required for intrinsic and optically evoked plasticity in D1 MSNs.

**Figure 3:**
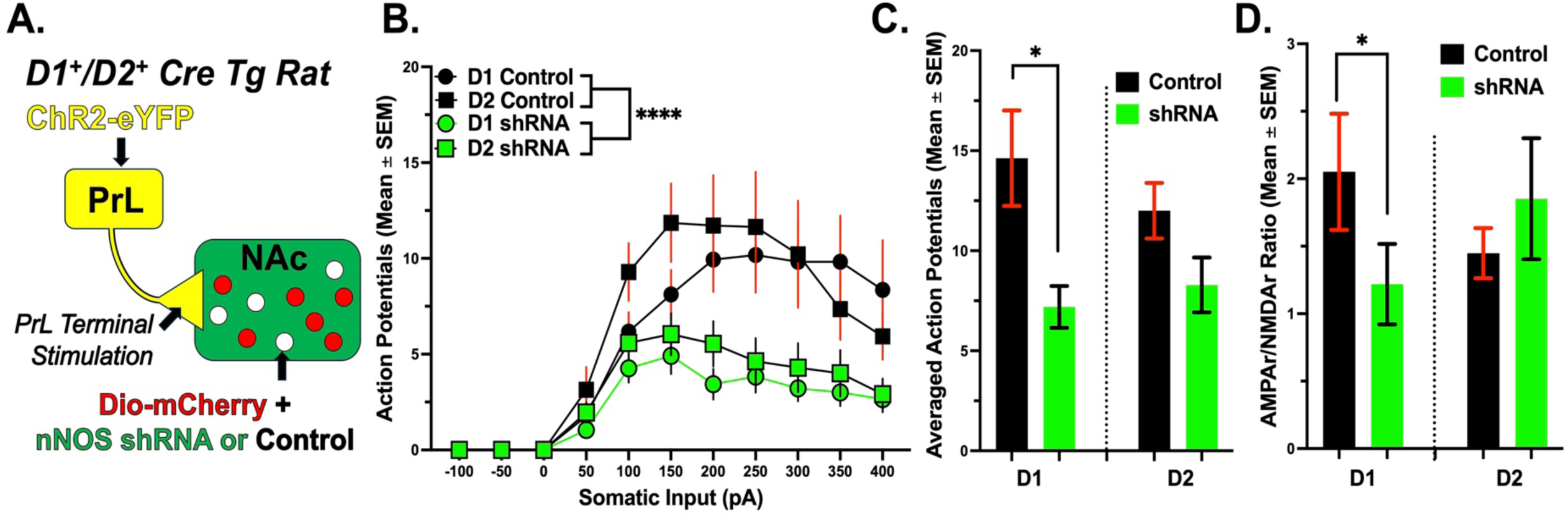
nNOS shRNA Reduces Intrinsic Excitability and AMPAr/NMDAr ratios in Cocaine Naïve Animals: **A.)** Schematic indicating viral design for NAc PrL terminal stimulation and viral labeling of *D1+/D2+* MSNs. **B.)** Averaged number of action potentials fired at increasing steps of injected current (-100-400pA at 50pA step increments). nNOS shRNA reduced the average number of action potentials in both *D1+* and *D2+* MSNs across stimulations. **C.)** Average maximum spike number for control and shRNA treated animals. nNOS shRNA significantly reduced average maximum spike numbers in *D1+* MSNs. While there was a marginal reduction for *D2+* MSNs, the effect was not statistically significant. **D.)** AMPAr/NMDAr ratios for control and nNOS shRNA treated animals. nNOS shRNA significantly reduced AMPAr/NMDAr ratios of *D1+* MSNs following PrL terminal stimulation in the NAc. While a marginal increase in AMPAr/NMDAr ratios was observed for *D2+* MSNs, the effect was not statistically significant. *\*p<0.05; ****p<0.001*.

### Experiment 2: nNOS is Required for Cocaine Cue-Induced D1 MSN Structural Plasticity

To evaluate the impact of knockdown of nNOS upon MSN spine head diameter (d_H_) adaptations associated with cued cocaine seeking, *D1* or *D2-Cre* transgenic rats were rapidly sacrificed immediately after 15 minutes of cued cocaine seeking to capture transient structural plasticity as described previously^6^. Next, tissue sections (3 per animal) were processed with mouse anti-Flag (Cre-dependent neuronal label indicating D1 or D2 MSNs). Isolated NAc MSN dendrites (∼5 per animal) were imaged, and 3D renderings were compiled using the “filaments” tool in Imaris. **Figure 4A** shows a schematic for viral labeling of NAc MSNs, while heatmaps of the localization of viral transduction are shown in **Figure 4B**. Nested ANOVAs were used to evaluate genotype, sex, drug-treatment, and viral treatment upon MSN head diameter (d_H_, **Figure 4C).** A main effect of cocaine SA, EXT and RST was observed for spine d_H_, indicating that cued RST increased the average spine d_H_ on NAc MSN’s [F(1, 58.31=13.41, p<0.001). Interestingly, a significant interaction effect between drug treatment and genotype emerged, indicating that increases in spine dH following cocaine RST were preferentially driven by D1 MSNs [F(1,58.31)=5.266, p=0.025]. A main effect of shRNA treatment was also observed, with nNOS shRNA treatment significantly reducing spine d_H_ overall, regardless of genotype and drug treatment, suggesting a role for nNOS in MSN structural plasticity that mirrors our findings with respect to functional plasticity. [F(1, 58.31)=6.647, p=0.012]. An interaction between viral treatment and genotype was likewise observed, with nNOS shRNA significantly reducing spine d_H_ in preferentially D1 but not D2 MSNs [F(1,58.31)=5.563, p=0.022], suggesting that nNOS-dependent structural plasticity adaptations following RST preferentially depend upon D1 MSNs. **Figures 4D-E** show representative co-expression of Flag and GFP (shRNA) and nNOS IHC. An isolated NAc MSN with selected dendritic segment is shown in **Figure 4F**. Binned spine d_H_ values separated by genotype are displayed in **Figure 5A-B**. Chi square (Ξ^2^) tests of independence indicated global distributional shifts in spine d_H_ for both D1 [Ξ^2^(12)=314.9, *p<0.001*] and D2 MSNs [Ξ^2^(12)=125.1, *p<0.001*]. Z-tests of proportions were then used to examine intra-bin differences between treatment groups. For D1 MSNs, there was a global trend of cocaine to increase frequency of large d_H_ spines, indicating that cocaine RST drives the formation of large spines, which was prevented by nNOS knockdown. Interestingly, knockdown of nNOS in yoked saline controls resulted in an increased frequency of small to medium spines, but a decreased frequency of large spines. While the overall Chi square was significant for frequency distributions of D2 d_H,_ follow-up tests reveal no treatment-dependent specificity in these changes. These results suggest that nNOS is required for cocaine cue-induced MSN structural plasticity and moreover indicate that structural plasticity adaptations following RST to cocaine are preferentially dependent upon D1 MSNs, with nNOS mechanistically linked to the cell-type specific spine d_H_ adaptations associated with cued cocaine seeking. Representative individual filaments selected for analysis are shown in **Figure 5C**. SA and active lever responding during EXT and RST for these animals is shown within **Figure 2D**

**Figure 4:**
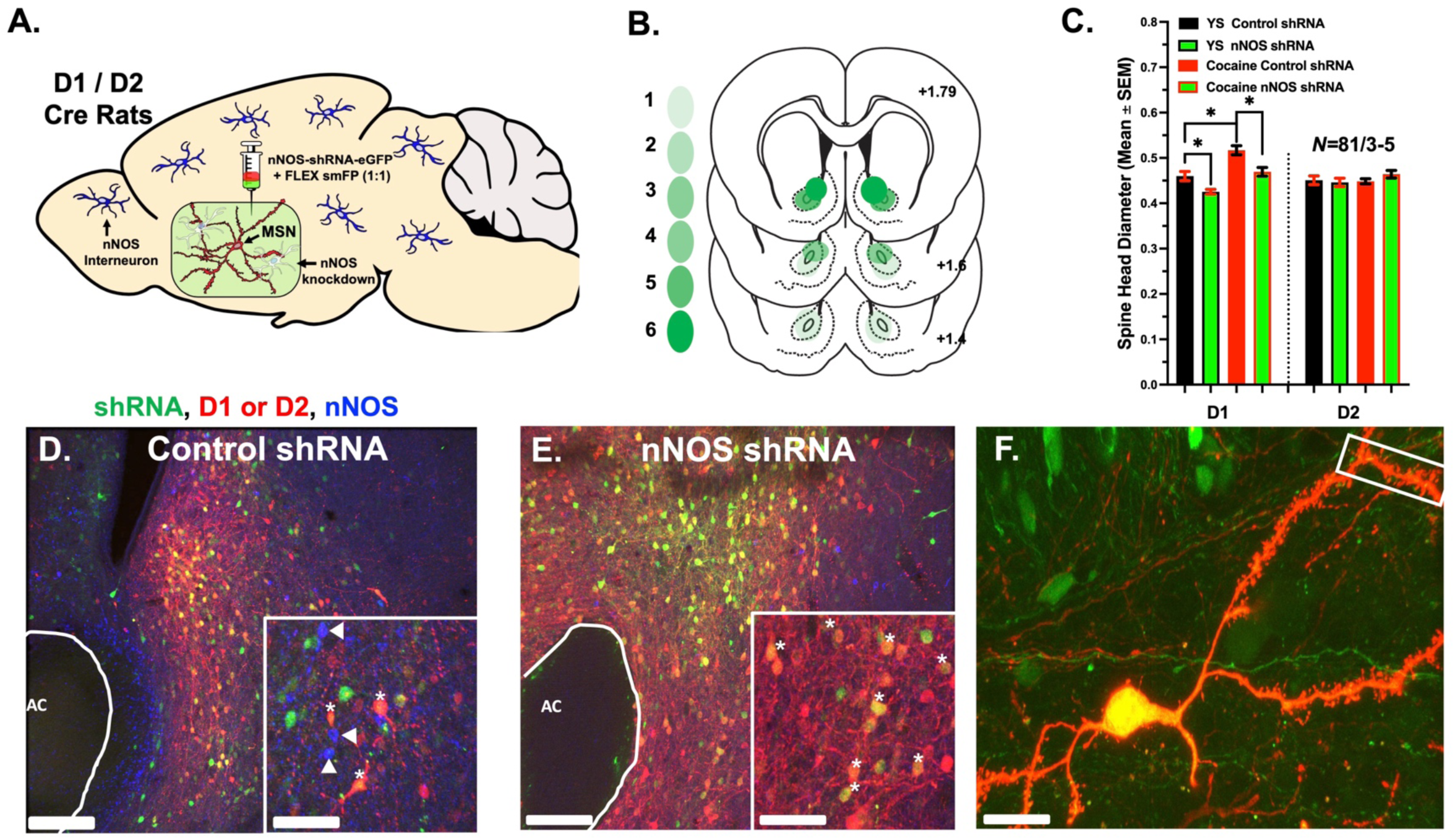
nNOS shRNA Inhibits *D1+* MSN Spine Head Expansion Associated with Cued Relapse to Cocaine: **A.)** Schematic indicating design for shRNA manipulation and viral labeling of *D1+/D2+* MSNs. **B.)** Heatmaps indicating viral placements in the NAc. **C.)** Spine head diameter for all treatment groups. Data are collapsed across sex as no sex effects were found. Cocaine RST significantly increased spine d_H_ relative to YS controls, while nNOS shRNA blocked increased spine d_H_ in *D1+* but not *D2+* MSNs. In cocaine naïve animals, nNOS shRNA resulted in a decreased spine d_H_ relative to control shRNA. **D-E.)** Representative images showing expression of EGFP (shRNA construct) Flag (*D1+/D2+* MSNs) and nNOS IHC. *Scale bars indicate 100 μm (low mag) or 50 μm (high mag) nNOS+ cells are indicated by triangles.* Asterisks indicate colocalization of EGFP (shRNA) and Flag (*D1+/D2+* MSNs). **F.)** Representative high magnification image of MSN chosen for analysis. Isolated dendritic segment chosen for analysis is indicated by white box; *Scale bar indicates 15 μm*. *\*p<0.05*.

**Figure 5:**
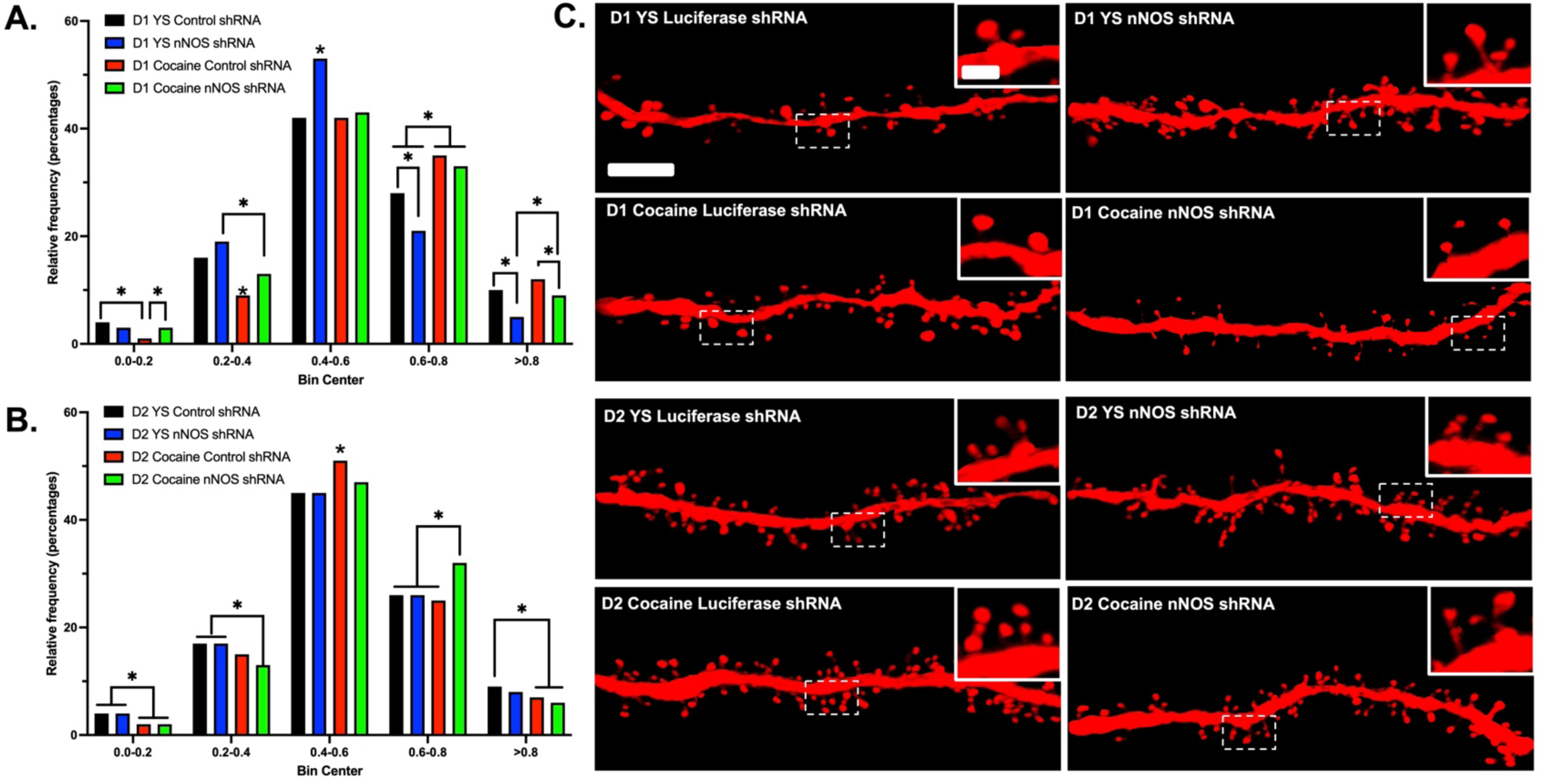
Frequency Distributions for Spine d_H_. **A.)** Frequency distributions of spine d_H_ for *D1+* MSNs. In YS animals, nNOS shRNA increased the frequency of smaller head diameters, while reducing the frequency of larger spine head diameters. Cocaine RST increased the frequency of larger head diameters relative to YS controls, and nNOS shRNA decreased the frequency of large diameter spine heads in cocaine reinstated animals. **B.)** Frequency distributions of spine d_H_ for *D2+* MSNs. Marginal differences in spine head diameter were observed across bin centers, though no global trends of treatment differences emerged. **C.)** Representative images for isolated dendritic segments for each treatment condition. *Scale bars indicate 3 μm (low mag) or 0.4 μm (high mag); *p<0.05*.

### Experiment 3-4: Selective Knockdown of mGluR5 in NAc Nitrergic Interneurons

Given our previous amperometry data suggested that application of mGluR5 agonists in anesthetized animlas can promote relese of NO in-vivo^2^, we hypothesized that mGluR5 receptors on NO interneurons are required for nNOS activation and NO release during cued cocaine seeking. To test this hypothesis we developed a Cre-dependent mGluR5 shRNA construct (**Figure 6A)** to use in concert with an inducible NOS1 Cre transgenic mouse line. Using our viral vector and transgenic animals, a subset of NosCre^ERT2^ mice were sacrificed and RNAScope™ was performed on NAcore sections for Nos1 and Grm5 in a subset of Luciferase (Control) shRNA (n=4) and mGluR5 (Grm5) shRNA (n=4) mice. As shown in **Figure 6B**, a nested t-test indicated that following administration of tamoxifen, NosCre^ERT2^ mice that received the Cre-dependent Grm5-shRNA construct showed an ∼80 reduction in Grm5 mRNA, selectively in nitrergic interneurons, within the zone of viral transduction relative to NosCre^ERT2^ mice that received the Cre-dependent Luciferase (control)-shRNA [t(41)=13.27, *p*<0.001]. **Figure 6C** shows high magnification images of isolated nitrergic interneurons (indicated by red Nos1 RNAscope puncta) and selective elimination of mGluR5 (blue Grm5 RNAscope puncta) on these interneurons.

**Figure 6:**
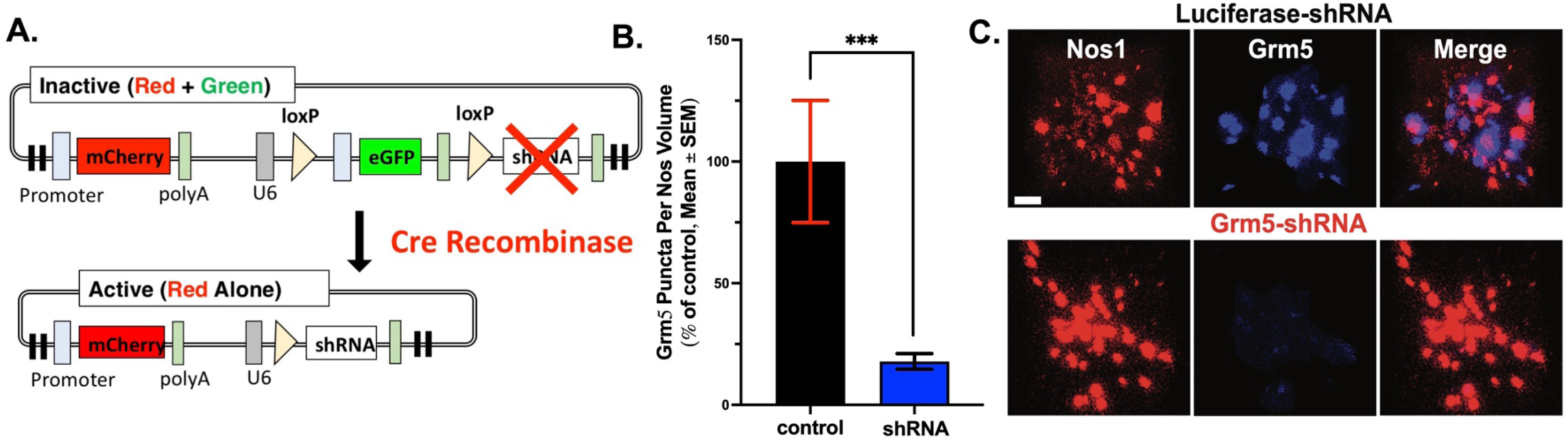
Validation of Novel mGluR5 (Grm5) shRNA construct. **A.)** Schematic of novel mGluR5 shRNA construct. **B.)** RNAscope quantification of mGluR5 puncta. mGluR5 shRNA reduced expression of mGluR5(Grm5) puncta by -90%. **C.)** Representative images of RNAscope experiment showing colocalization of Nos1 and Grm5 puncta (control) and selective lack of Grm5 colocalization with Nos1 puncta (shRNA). *Scale bars indicate 3 μm; ***p<0.01*.

### Experiment 3 Expression of mGluR5 on NO Interneurons is Required for Cocaine CPP

To investigate the role of mGluR5 receptors expressed on NAc nitrergic interneurons in cocaine-related behaviors, we first examined whether these receptors are required for cocaine conditioned place preference (CPP). **Figures 7A-B** shows the surgical schematic for viral implantation and heatmaps of viral expression. A schematic for the design of our CPP experiment is shown in **Figure 7C**. As shown in **Figure 7D**, selective mGluR5 knockdown from NAc NO interneurons prior to cocaine CPP conditioning suppressed the expression of cocaine place preference during the post-test, confirming the requirement of mGluR5 on nitrergic interneurons [Welch’s t (21.75)=2.46, *p*=0.022; data collapsed across sex] in cocaine associative reward.

**Figure 7:**
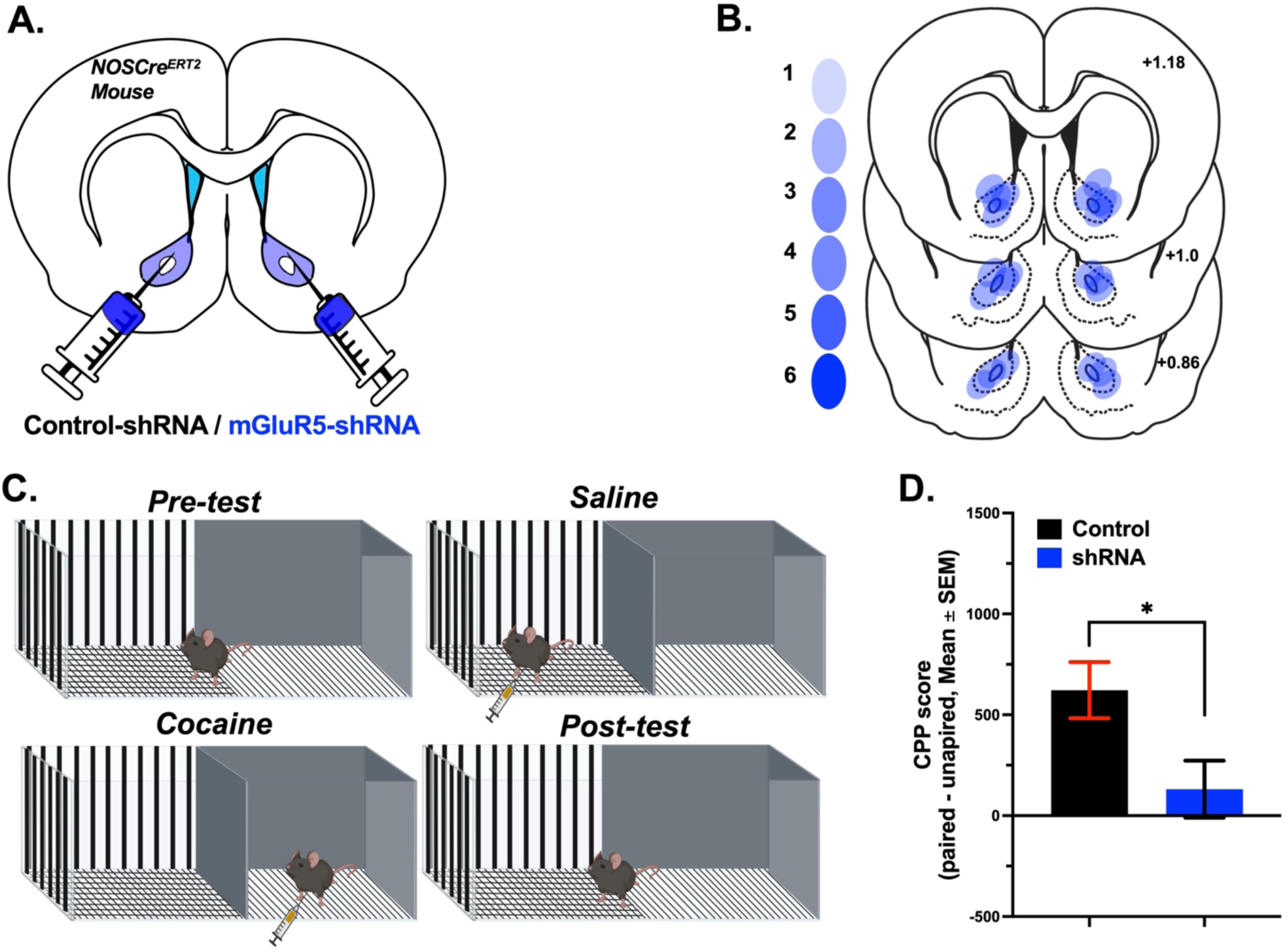
mGluR5 shRNA Blocks Cocaine CPP. **A.)** Schematic indicating surgery design of CPP experiment. **B.)** Heatmaps indicating viral placements in the NAc. **C.)** Schematic indicating experimental design of CPP experiment. **C.** CPP scores upon post-test. mGluR5 shRNA resulting in a statistically significant reduction of cocaine CPP. *\*p<0.05*.

### Experiment 4: Expression of mGluR5 on NO Interneurons is Required for Cued Cocaine Seeking

To determine if expression of mGluR5 on nitrergic interneurons is required for cued cocaine seeking we implanted our Cre dependent mGluR5 shRNA construct or a Cre-dependent control into a separate cohort of NosCre^ERT2^ mice (**Figure 8A**). Heatmaps of viral expression are shown in **Figure 8B**. **Figure 8C** indicates a timeline for behavioral training. Following SA, we administered 20 mg/kg 4-OHT. Following 4-OHT administration mice were given one week of homecage abstinence to allow for viral incubation. As expected, viral treatment (Cre dependent mGluR5 shRNA vs. control) was not found to alter SA acquisition or extinction, and both treatment groups demonstrated significantly higher ALP compared with ILP, indicating acquisition of behavior [F(1,13)=81.56, *p<0.001;* **Figure 8D**]. No differences were observed with respect to EXT. Knockdown of mGluR5 on NO interneurons blocked cue induced reinstatement to cocaine [F(1,11)=5.55, *p=0.04;* **Figure 8E**].

**Figure 8:**
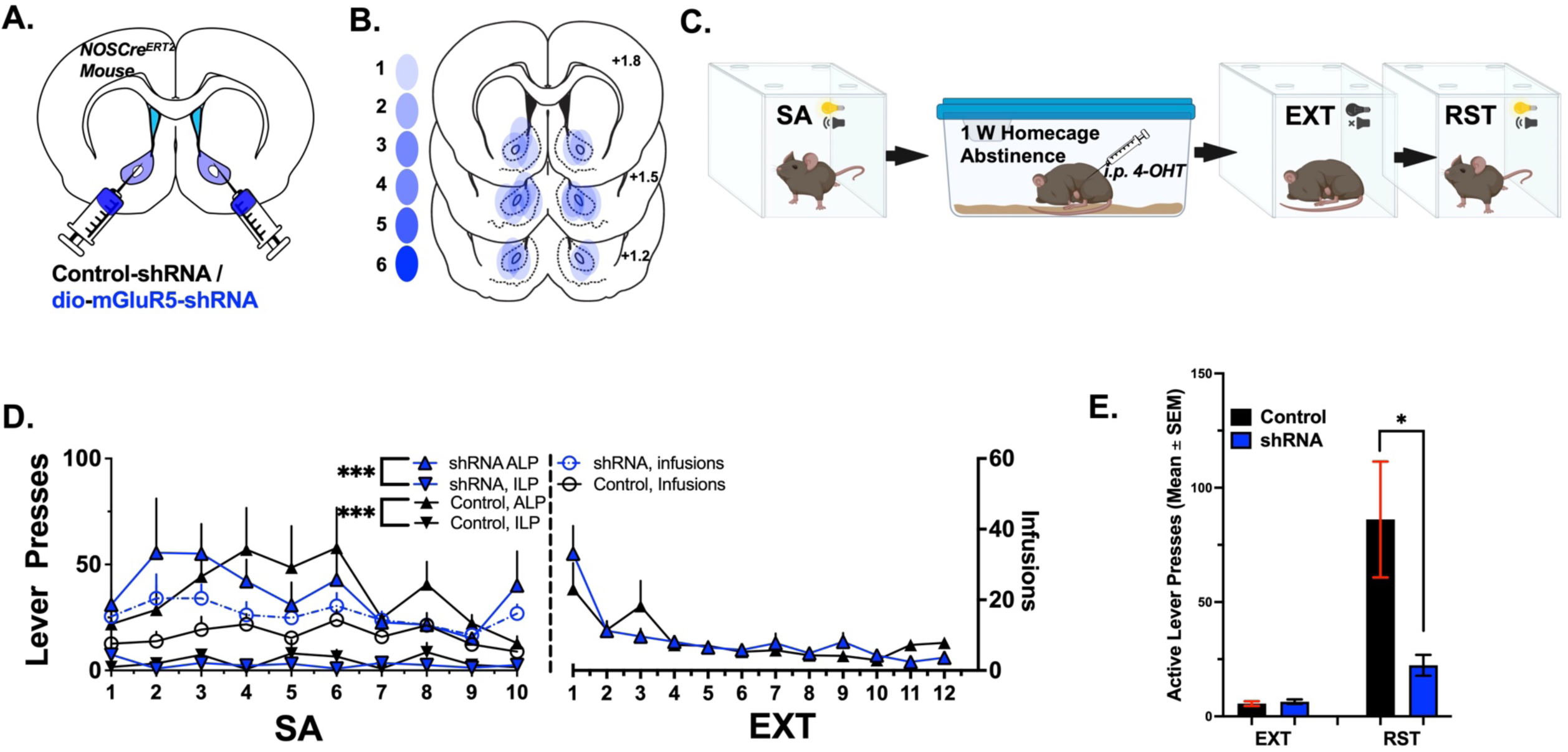
mGluR5 shRNA Blocks Cued Relapse to Cocaine. **A.)** Schematic indicating surgery design of experiment. **B.)** Heatmaps indicating viral placements in the NAc. **C.)** Schematic indicating experimental design for behavioral experiments. **D.)** SA and EXT data for all animals. Data are collapsed across sex as no sex differences were observed. No differences between viral treatment (control vs. shRNA) were observed across behavioral testing. Within each treatment condition, mice significantly discriminated between the active and inactive lever, indicating acquisition of SA behavior and motivated lever pressing. **E.)** Cued reinstatement data for control and shRNA treated animals. mGluR5 shRNA significantly reduced active lever responding during cued reinstatement testing. No sex differences were found during cued reinstatement testing. *\*p<0.05; ***p<0.001*.

### Experiment 5

Previous reports indicate that MMP activation underlying cued cocaine seeking engages β3 integrin activation which underlies MSN plasticity required for relapse ^36^, and that β3 integrin-mediated cofilin signaling was preferentially increased on D1 MSNs^32^. Given the primary involvement of D1 MSNs underlying cued cocaine seeking ^12,13^, we hypothesized that nNOS/NO dependent modifications to D1 MSNs depends on β3 integrin signaling. We implanted a Cre-dependent shRNA targeted to β3 integrin (**Figure 9A**) into D1/D2^Cre+^ rats and trained the animals to self-administer cocaine (**Figure 9B**). SA and EXT data are collapsed across genotype as no differences in behavioral responding were observed. Upon RST testing (**Figure 9C**), β3 integrin inhibition on D1 MSNs significantly reduced cued relapse responding, while inhibition on D2 MSNs did not [F(2,28)=4.91, *p=0.01,* Tukey MCT= 0.004 (control shRNA vs. D1 β3 shRNA), 0.003 D1 β3 shRNA vs D2 β3 shRNA. **Figure 9D** shows knockdown of β3 integrin for control (left) and shRNA (right) viral conditions.

**Figure 9:**
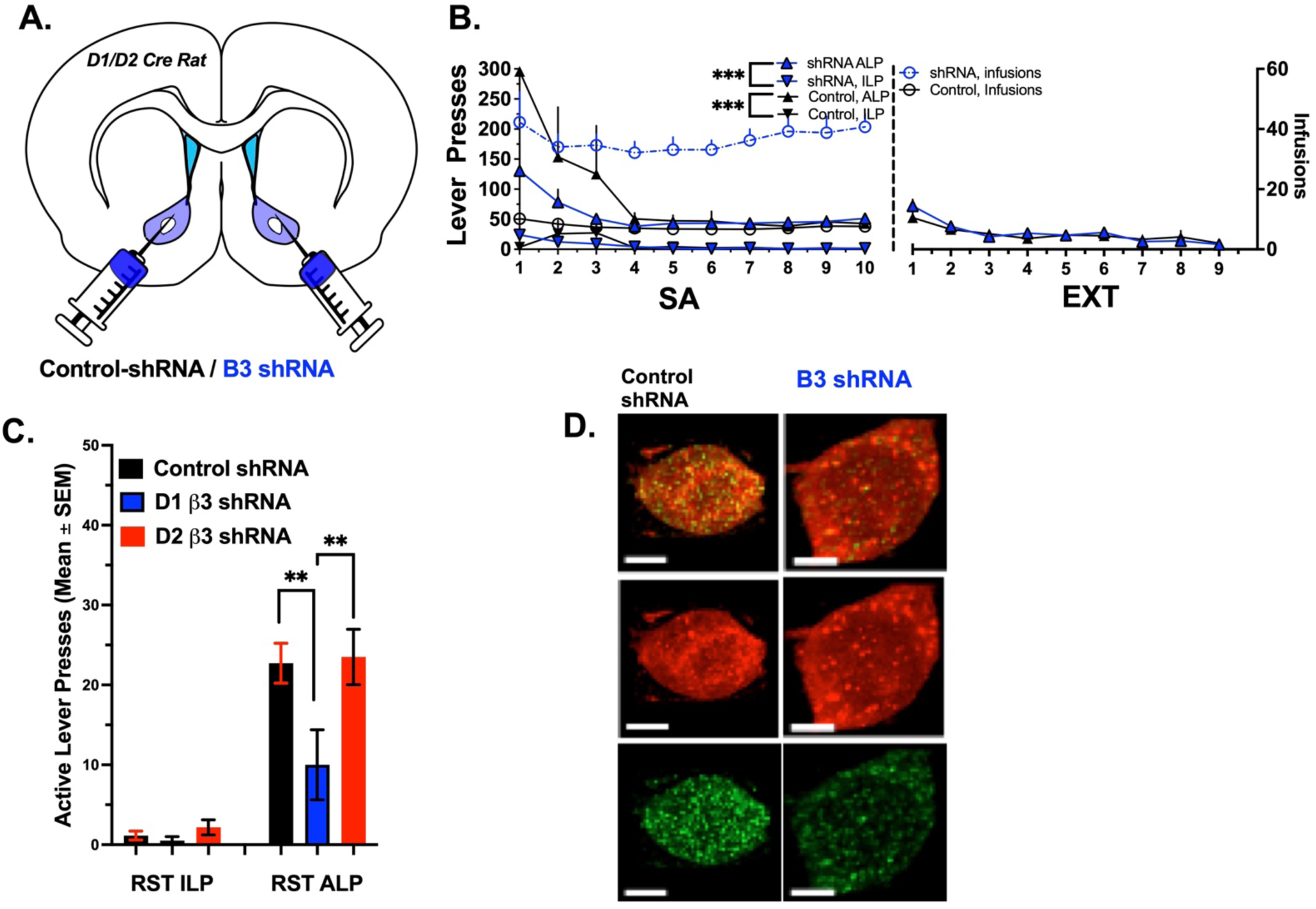
D1 β3 Integrin is Required for Cued Cocaine Seeking. **A.)** Schematic indicating surgery design. **B.)** SA and EXT data for all experiments. No differences were observed between rodent genotype or shRNA treatment across SA or EXT, and all animals successfully demonstrated behavioral acquisition and lever discrimination. **C.)** Cued relapse data. D1 β3 shRNA reduced ALP responding compared with both control shRNA and D2 β3 shRNA. **D.)** Representative images of D1 (mCherry tag) colocalization with β3 integrin for control shRNA (left) and β3 shRNA (right). *\*\*p<0.01*

## Discussion

Here, we show that RST to cocaine seeking following EXT training requires the activation of nNOS. Knockdown of nNOS in the NAc via NOS1 shRNA significantly reduced the number of nNOS+ cells evaluated by immunohistochemistry, and blunted RST to cocaine following EXT training, a finding which adds a needed confirmation of our previous data that used correlation analyses and selective death of nNOS^+^ neurons via a Cre-dependent caspase based strategy^2,11^. Moreover, we reproduce the finding that D1 MSNs predominantly drive structural plasticity following RST to cocaine^12,13^. Critically, knockdown of nNOS blocks structural plasticity adaptations during cocaine RST, selectively in D1 MSNs but does not impact spine head d_H_ in D2 MSNs. Further, knockdown of nNOS impairs intrinsic excitability and PrL input-dependent optically evoked AMPA/NMDA ratios on D1 MSNs. We likewise show that expression of mGluR5 on NAc NO interneurons is required for both cocaine CPP and cued-cocaine seeking, and finally demonstrate that expression of β3 integrin receptors on D1 MSNs, which we have previously shown are critical mediators of NO-induced MMP turnover^2,32,36^ is likewise required for cued cocaine seeking. Collectively, our data provide mechanistic insight into the requirement of nitrergic interneurons as a central node in the signal transduction cascade within the NAc that translates cortically derived cocaine cue-induced glutamate release in the NAc into the D1 MSN plasticity required for cocaine seeking.

Rodent models of rodent SA, EXT, and relapse indicate that SA and extinction to cocaine, and other drugs of abuse, engages maladaptive glutamatergic homeostasis in the NAc. ^6,10,17^. Specifically, cocaine exposure reduces astrocyte derived basal extrasynaptic glutamate levels in the NAc, leading to presynaptic potentiation due to reduced autoreceptor activation^37^. In concert with this maladaptive response, cocaine-mediated disruptions in the ability of astrocytes to clear synaptically released glutamate give rise to an accumulation of glutamate in the NAc, which is required to drive cocaine seeking^9,10,19,38–40^. We have previously demonstrated that the major source of glutamate release in the NAc during cue-induced cocaine seeking is the PrL, with inhibition of this pathway bringing NAc glutamate levels to those observed during cued sucrose seeking^16^. Importantly, normalization of NAc glutamate homeostasis with compounds such as N-acetylcysteine or ceftriaxone prevents cue-induced drug seeking^39,41^.

Downstream of cue-induced glutamate release, cue-induced cocaine seeking requires rapid transient plasticity in MSNs. Specifically, cue-induced glutamate release engages enlarged spine head diameter in existing dendritic spines and plasticity in MSNs, with the magnitude of these adaptations being positively correlated with the magnitude of relapse responding ^6,16,42^. Importantly, cue-dependent synapse remodeling requires the degradation of extracellular matrix proteins via matrix metalloproteinases (MMPs) 2 and 9 ^30,43^. Pharmacological inhibition of MMPs 2 and 9 has been demonstrated to prevent LTP increases in MSNs as measured by AMPA/NMDA ratios, and it has moreover been demonstrated that spine d_H_ increases during extinction and cued-cocaine seeking are likewise dependent upon MMPs -2 and 9, respectively.

The NAc is composed mainly of projecting MSNs accompanied by a sparse population of interneurons. NAc MSNs preferentially innervate the dorsolateral ventral pallidum, the medial substantia nigra, and the ventral mesencephalon^44–46^. MSNs can further be subdivided into D1 or D2 receptor expressing MSNs^47^. D1 MSNs projecting from the NAc positively regulate motivational action via disinhibition of connected basal ganglia structures, comprising a “direct” pathway that is hypothesized to promote motivated behavior^46,48^. Recent studies have indicated that the ventral pallidum is the primary target for D1 MSNs in the NAc that motivate relapse to cocaine, with inhibition of this pathway blocking cue-induced reinstatement^46^. D2 MSNs project broadly throughout the basal ganglia and serve to inhibit the function of downstream brain regions, providing negative regulation of motivational states ^46,48,49^. This proposed dichotomy of striatal MSNs gave rise to the “classic” or “go/no go” model of striatal MSN function^50,51^.

This “classic” model posited two major classifications of striatal MSNs into direct (dMSNs) or indirect (iMSNs). dMSNs were observed to project directly to the midbrain while iMSNs were observed to project indirectly to the midbrain through the pallidus and subthalamic nucleus ^50,52–54^. Early studies of dopamine and locomotion showed that dMSNs predominantly express the D1 receptor while, iMSNs predominantly express the D2 receptor. As an important note, however, much of this early work was performed in the dorsal striatum as opposed to the ventral striatum, and more recent work has suggested that this classic dichotomy may not be appropriate to characterize the ventral striatum/NAc^49,51^. An increasing body of research indicates important functional and structural adaptations in D1 MSNs specifically following cocaine exposure^42,55–58^. Studies employing fiber photometry have reported that activation of D1 MSNs is an essential requirement for cocaine-context associations and for relapse to drug seeking following extinction in a conditioned place preference model of cocaine exposure^59^. Importantly, relapse associated plasticity adaptations and MMP activity have been directly linked to D1 MSNs, with the majority of neuronal ensembles recruited during cocaine seeking being comprised of D1 MSNs ^13^ and spine d_H_ enlargement driven largely by D1 MSNs ^32^. In studies of cued heroin seeking, MMP-9 activity has been demonstrated to be increased preferentially around D1-MSNs ^60^. With respect to cocaine seeking, MMP-9 activation stimulates β3 integrin signaling through focal adhesion kinase (FAK) activation to promote both cued-seeking and MSN plasticity^36^. More recent investigations demonstrated that β3 integrin-mediated FAK activation was increased in both D1 and D2 MSNs, but β3 integrin-mediated cofilin activation was preferentially increased in D1-MSNs, consistent with previous findings indicating that relapse-associated spine plasticity is driven by D1 MSNs ^12,32^.

The role of nNOS and NO in the signaling cascade described above has until relatively recently been unclear, with few studies directly investigating the role of NO in cued cocaine seeking in the context of D1 vs. D2 MSNs. NO in the NAc is produced by a subpopulation of interneurons comprising <1% of the total cellular composition of the NAc, with Ca^2+^ -dependent nNOS required for NO production ^2,11^. As discussed above, altered glutamatergic homeostasis in the NAc following cocaine withdrawal is associated with reduced activation of presynaptic auto receptors like mGluR2/3, with activation of these receptors inhibiting cued cocaine seeking^27^. An opposing effect is observed with respect to mGluR5 receptors, however, with *blockade* of mGluR5 receptors inhibiting cued cocaine seeking ^28,61^. In an elegant series of experiments, Smith and colleagues demonstrated that pharmacological activation of mGluR2/3 prevented cue-induced increases in extracellular glutamate concentrations ^2^, while blockade of mGluR5 did not prevent glutamate accumulation but does prevent cued cocaine seeking. Gq-coupled mGluR5 receptors promote the release of calcium from intracellular stores and are expressed on most NAc neurons, including NO interneurons.^62^ Given that nNOS activation is calcium-dependent ^63^ and consequent NO release regulates synaptic plasticity ^64^, Smith and colleagues reasoned that mGluR5 activation during cued cocaine seeking may have direct impact on NO signaling^2^. Indeed, they showed that pharmacological stimulation of mGluR5 with CHPG induced a dose-dependent increase in NO release in the NAc and microinjection with CHPG increased active lever responding in the absence of cocaine conditioned cues^2^. Moreover, using NPLA, an NO inhibitor, Smith et al., demonstrated that pharmacological inhibition of NO blocked both cued seeking and associated MMP activation^2^. In our recent work we have mapped the time-course of glutamate and NO release, with amperometric recordings demonstrating that peak glutamate and NO release occur within 15-20 minutes of cocaine cue exposure, with NO levels detected approximately 1-2 minutes later than glutamate^11^, a time-course which parallels the peak of associated plasticity in MSNs ^6^. Downstream of this glutamate dependent signaling, it has also been demonstrated that nNOS/NO activity is required for MMP-2 and MMP-9 activation during extinction and cued seeking, respectively, with nNOS inhibition reducing MMP activity at these respective time points^2^. In the same study using NOS1-Cre mice, it was demonstrated that Gq-DREADD activation of NO interneurons in drug naïve mice increased AMPA/NMDA ratios in NAc MSNs and that similar stimulation was sufficient to recapitulate cocaine seeking in the absence of cocaine conditioned cues ^2^.

Here, we demonstrate that nNOS/NO is directly required for cued cocaine seeking, showing that shRNA-mediated knockdown of nNOS blocked cued-reinstatement to cocaine. Further^2^, in drug naïve animals nNOS knockdown reduced AMPA/NMDA ratios upon PrL terminal stimulation in preferentially D1 MSNs, consistent with previous reports showing activation of NO increased AMPA/NMDA ratios in MSNs^30^, as we show that inhibition of nNOS/NO reduced MSN AMPA/NMDA ratios and further show that this reduction is cell-type specific. Replicating previous findings ^12,13,32^, we show here that structural plasticity in MSNs is likewise cell-type specific and driven by predominantly D1-MSNs. Importantly, we show that nNOS is required for D1 MSN plasticity, as nNOS knockdown prevented cue-associated increases in D1 MSN d_H_.

We also demonstrate that knockdown of mGluR5 on NO interneurons blocks cocaine CPP and cued cocaine seeking. These data support our overall model of glutamate-mediated activation on nNOS neurons in the molecular mechanism underlying cocaine seeking behavior. Finally, we demonstrate that knockdown of β3 integrin on D1 MSNs is sufficient to block cued cocaine seeking, supporting an ECM-linked mechanism through which cortically derived cue-induced glutamate release is translated into cell type-specific MSN plasticity. (**Figure 10**).

**Figure 10:**
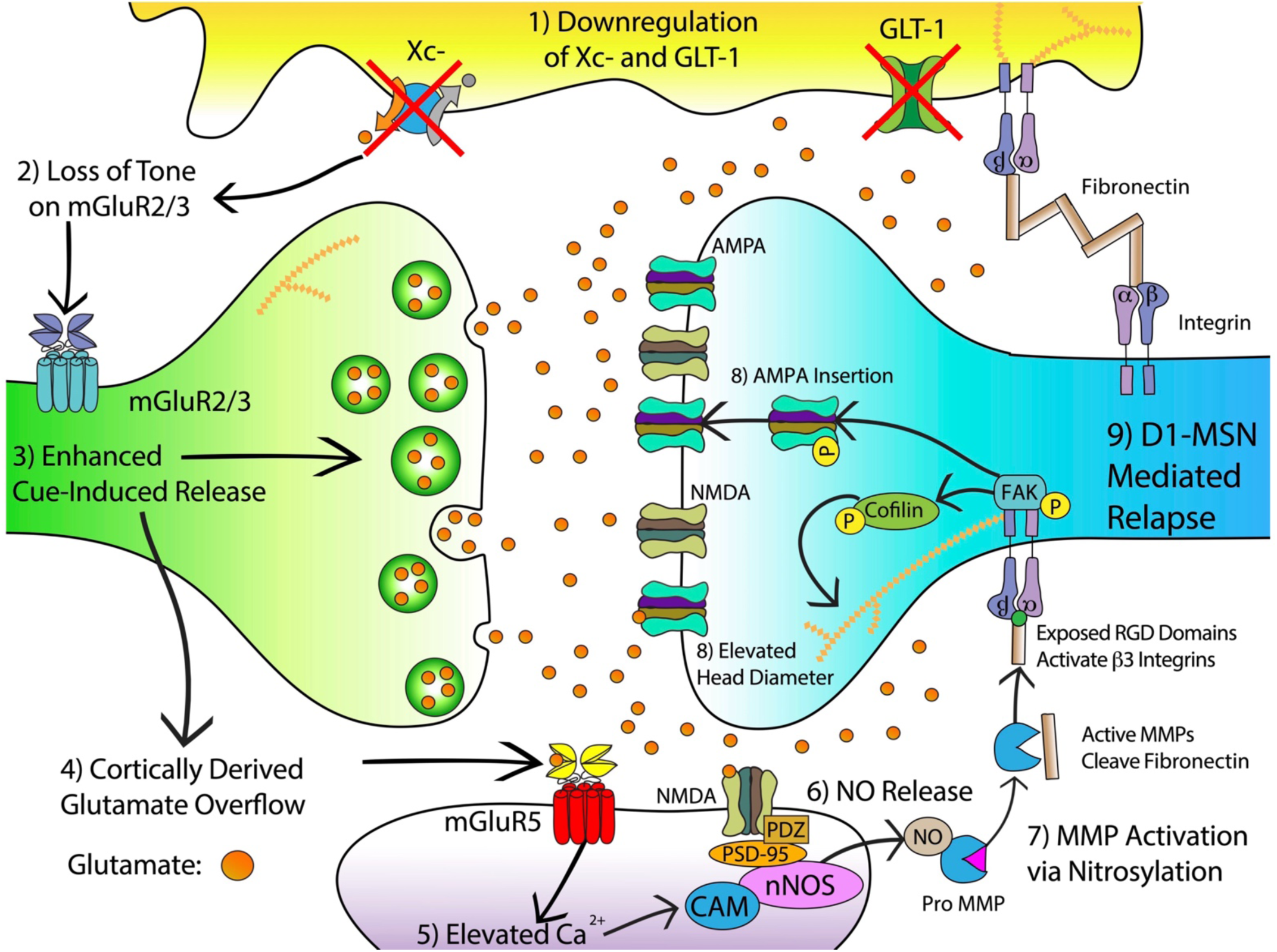
Summary Figure of Relationship Between Cue-Evoked Glutamate, NO and Structural/Functional Plasticity in D1 MSNs.

## Notes

### Competing Interest Statement

The authors have declared no competing interest.

